# The yeast eIF2 kinase Gcn2 facilitates H_2_O_2_-mediated feedback inhibition of both protein synthesis and ER oxidative folding during recombinant protein production

**DOI:** 10.1101/2021.02.03.429681

**Authors:** Veronica Gast, Kate Campbell, Cecilia Picazo Campos, Martin Engqvist, Verena Siewers, Mikael Molin

## Abstract

Recombinant protein production is a known source of oxidative stress. Knowledge of which ROS are involved or the specific growth phase in which stress occurs however remains lacking. Using modern, hypersensitive genetic H_2_O_2_-specific probes, micro-cultivation and continuous measurements in batch culture, we observed H_2_O_2_ accumulation during and following the diauxic shift in engineered *Saccharomyces cerevisiae*, correlating with peak α-amylase production. In agreement with previous studies supporting a role of the translation initiation factor kinase Gcn2 in the response to H_2_O_2_, we find Gcn2-dependent phosphorylation of eIF2α to increase alongside translational attenuation in strains engineered to produce large amounts of α-amylase. Gcn2 removal significantly improved α-amylase production in two previously optimized high-producing strains, but not in the wild-type. Gcn2-deficiency furthermore reduced intracellular H_2_O_2_ levels and the unfolded protein response whilst expression of antioxidants and the ER disulfide isomerase *PDI1* increased. These results suggest protein synthesis and ER oxidative folding to be coupled and subject to feedback inhibition by H_2_O_2_.

**Importance:** Reactive oxygen species (ROS) accumulate during recombinant protein production both in yeast and Chinese hamster ovary cells, two of the most popular organisms used in the multi-million dollar protein production industry. Here we document increased H_2_O_2_ in the cytosol of yeast cells producing α-amylase. Since H_2_O_2_ predominantly targets the protein synthesis machinery and activates the translation initiation factor kinase Gcn2, we removed Gcn2, resulting in increased recombinant α-amylase production in two different previously engineered high-producing protein production strains. Removal of this negative feed-back loop thus represents a complementary strategy for improving recombinant protein production efforts currently used in yeast. Gcn2-deficiency also increased the expression of antioxidant genes and the ER-foldase *PDI1*, suggesting that protein synthesis and ER oxidative folding are linked and feed-back regulated via H_2_O_2_. Identification of additional components in this complex regulation may further improve protein production and contribute to the development of novel protein-based therapeutic strategies.

## Introduction

The biotechnological role of *S. cerevisiae* in the production of bread and beer has been long established. In recent decades however, this yeast has also proven effective as a host for the production of recombinant proteins of significant pharmaceutical value (1,2). *S. cerevisiae* is a successful production host predominantly due to its eukaryotic post-translational modification machinery, its ability to secrete proteins to the media, as well as its robustness to harsh industrial conditions amongst other traits (2, 3). Many different strategies have been shown to improve recombinant protein production and secretion in yeast (4, 5) including the engineering of transport mechanisms in the secretory pathway, increasing the expression of chaperones, as well as even expanding the size of the endoplasmic reticulum (ER) (6–9).

Recombinant protein production is, however, known to be a significant burden for cells, due to for example limiting secretory capacity and protein misfolding (10). In engineered high-producing strains in particular, this burden is speculated to increase concomitantly with production levels, leading to ER stress (11, 12). To counter this and the accumulation of unfolded proteins within this organelle, two response mechanisms can be activated; the unfolded protein response (UPR) and ER-associated degradation (ERAD). The UPR in *S. cerevisiae* is initiated by Ire1, an ER membrane protein with active subunits both in the ER lumen and on the cytosolic side. Upon Ire1 activation by ER stress, an mRNA encoding a transcription factor, Hac1, is spliced to its active form. Hac1p subsequently moves to the nucleus and activates the expression of UPR-associated genes (13)

Besides organelle-specific stress response mechanisms, eukaryotic cells also mount the general stress response. An example of this is the phosphorylation of the α-subunit of the eIF2 translation initiator factor (eIF2α) (14), which leads to the attenuation of general translation and a reduction in protein synthesis. Mammals have a total of four kinases that can phosphorylate eIF2α in response to various stress signals, *PERK, PRK, GCN2*, and *HRI*, whereas *S. cerevisiae* only expresses one of these, *GCN2* (14). The protein kinase Gcn2 in *S. cerevisiae* is mainly known as the activator for the general amino acid control (15). Upon depletion of one or multiple amino acids, this response is activated to counteract amino acid depletion. Besides reducing translation, a downstream target of Gcn2 within the general amino acid control is the transcription factor Gcn4. Gcn4 is translationally regulated and activates the expression of genes involved in the biosynthesis of amino acids amongst other targets (16). However, over the years more conditions other than amino acid starvation have shown to activate Gcn2. As these stresses have also led to general translation attenuation, the Gcn2-mediated response has subsequently been renamed the integrated stress response (15–20).

One of the stresses known to activate the protein kinase Gcn2 in *S. cerevisiae* is H_2_O_2_ (21), a signaling molecule, as well as a byproduct of multiple biochemical reactions. Intracellular levels of H_2_O_2_ and other reactive oxygen species (ROS) are usually maintained below certain thresholds to avoid deleterious effects, such as untargeted oxidation of cellular components (DNA, lipids and protein) as well as in extreme cases, cell death (apoptosis) (22–24). When levels of ROS do exceed this threshold, cells are known to respond by upregulating anti-oxidant proteins, redirecting metabolism as well as attenuating growth responses such as the protein synthesis machinery to regain homeostasis (25).

Oxidative phosphorylation in the mitochondria and protein production in the ER can both be major sources of ROS (26, 27). Recombinant protein production also has shown to induce both ER stress and oxidative stress (26, 28). Within the ER, oxidative stress is suggested to arise due to H_2_O_2_ production during protein folding (11, 12). H_2_O_2_ is a direct byproduct of the reduction of oxygen, which occurs during disulfide bond formation, an iterative process mediated by Pdi1 and Ero1 (15). Oxidative stress subsequently limits protein secretion in both Chinese hamster ovary (CHO) cells and yeast (6, 26), with the production capacity of ‘super-producer’ engineered strains most likely experiencing this limitation as well.

We hypothesize that recombinant protein production could induce a negative feedback loop mediated by Gcn2 resulting in the reduction of translation and protein synthesis. In this study, we provide evidence for the production of H_2_O_2_ during recombinant protein production, using hypersensitive peroxiredoxin-based probes (29). Furthermore, by removing the H_2_O_2_-activated translational initiation factor kinase Gcn2 we were able to enhance recombinant α-amylase production in *S. cerevisiae* by 75%. We find improved recombinant protein production to also correlate with the induction of the disulfide isomerase encoding gene *PDI1* as well as several antioxidants and reduced H_2_O_2_ levels. Based on this data we propose a model in which protein synthesis and ER-folding are coupled and subject to feedback-inhibition via H_2_O_2_ and Gcn2.

## Results

### Recombinant α-amylase production leads to elevated levels of H_2_O_2_ in the engineered strain B184

Previous work has shown oxidant production to limit recombinant protein production and secretion in yeast and CHO cells respectively (6, 26). In both these studies, the fluorescent probes used to assess oxidant production suffered from low specificity, with their response to ROS levels being impacted by peroxidase activity as well as metal ion levels. Information on the specifics of oxidant production during protein secretion subsequently remains lacking (30). Recombinant protein productivity in batch cultivation is also speculated to differ across different growth phases. Measuring this necessitates oxidant production to be monitored continuously (6), enabling subtle changes in H_2_O_2_ to be identified during different phases of cell growth. To address this, we decided to use peroxiredoxin-linked redox-sensitive (ro) GFP sensors (29) in combination with micro-cultivation (31), considering that peroxiredoxins are by far the most H_2_O_2_-reactive proteins in the cell (32). Upon oxidation of the sensor, peroxiredoxin and redox-relay to the fused roGFP2, and a fluorescent signal excited at a wavelength of 405 nm is emitted by the sensor; upon sensor reduction this signal is instead excited at a wavelength of 488 nm. By calculating the ratio of oxidized to reduced signal (Ox/Red), we were able to compare the internal H_2_O_2_ levels in different strains. We initially started with three sensors, roGFP2-*Pf*AOP, roGFP2-*Pf*AOP^L109M^, and roGFP2-Prx1, and investigated their responses to external addition of H_2_O_2_ and DTT (Figure S1) (29, 33). We found the roGFP2-Prx1 sensor (Ox/Red) ratio to increase upon H_2_O_2_ addition and decrease upon DTT addition, whereas both the roGFP2-*Pf*AOP sensors responded mainly to DTT addition (Figure S1). Importantly, the growth of the strains expressing the roGFP2-Prx1 sensor was also similar to the wildtype (Figure S2). Based on these results, we continued our experiments only with the roGFP2-Prx1 sensor, considering that this sensor demonstrated a high sensitivity to endogenous H_2_O_2_ levels (responded to DTT), while its signal still increased upon addition of exogenous H_2_O_2_ (Figure S1). Within this setup, we also subtracted yeast cell autofluorescence from the fluorescent signal of the roGFP2-Prx1 sensor. This being possible due to our strains harboring the roGFP2-Prx1 sensor and the vector control plasmid respectively having highly similar growth profiles (Figure S3-S4 Using our selected sensor, we next sought to study the impact of different levels of recombinant protein production on ROS generation. Here, we made use of B184 and AACK strains, two commonly used strains for recombinant protein production purposes. AACK is the ‘wildtype’ to B184, a strain based on AACK, which has also been engineered by random UV-mutagenesis to produce a 6-fold higher α-amylase titer in batch bioreactors (34, 35). α-amylase is used biotechnologically to release fermentable sugars from starch and is a commonly used marker protein to report on the recombinant protein production capacity in yeast cells (5, 9). We tested both strains to determine if a difference in ROS production could be observed as a consequence of their difference in capacity for α-amylase production. Based on the determined (Ox/Red) ratios, we found that recombinant α-amylase production led to increased H_2_O_2_ levels in strain B184 relative to the non-producing strain, with this increase predominantly occurring in the later stages of growth (Figure 1A). Since B184 demonstrates higher α-amylase production compared to AACK, the difference in Ox/Red ratios observed may be related to the amount of recombinant protein produced (35). In particular, we observed elevated (Ox/Red) ratios from around 25 h to the end of 96 h cultivation in B184 with recombinant α-amylase production i.e., during and following the diauxic shift (Figure 1A). Furthermore, (Ox/Red) ratios levels exhibited a cell density-dependent pattern in both B184 strains which may be related to oxygen levels and/or growth phase, as previously observed (Figure S3) (29). In AACK, the difference with and without α-amylase production was less pronounced however with the apparent peak in the (Ox/Red) ratio between 30 h and 50 h most likely being the result of delayed growth (Figure 1B & S3).

**Figure 1.**
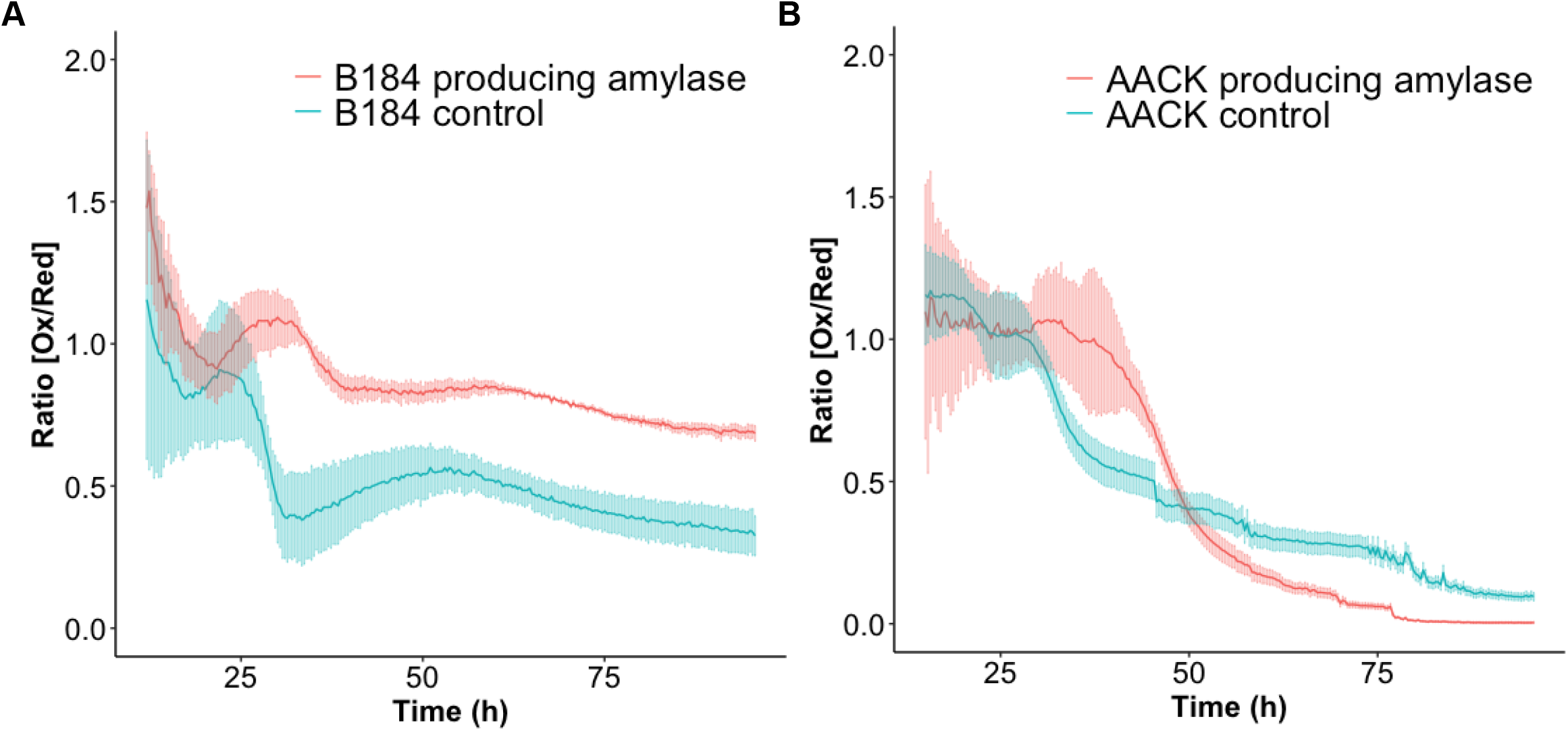
α-Amylase production leads to higher levels of intracellular H_2_O_2_ in the engineered high-level production strain B184. (Ox/Red) ratios over 96 h of cultivation for B184 and AACK measured with plasmid based *roGFP-PRX1*. B184 **(A)** and AACK **(B)** expressing α-amylase (red) and the control without expressing α-amylase (green). The light bars represent the standard deviations of three biological replicates and two technical replicates. The first 15 h were excluded due to too low signal.

### The protein kinase Gcn2 is active in B184 both with and without recombinant α-amylase production

Previous research suggests that external H_2_O_2_ addition activates the protein kinase Gcn2 leads to a reduction in protein synthesis (21), in part through its phosphorylation of the α subunit of the translation initiation factor (eIF2α). With the assumption that eIF2α would also respond to the increased H_2_O_2_ levels detected upon α-amylase production, we therefore monitored Gcn2-dependent phosphorylation of eIF2α in B184 and AACK ± α-amylase expression, by immunoblotting against total and phosphorylated eIF2α. Only B184 producing recombinant α-amylase showed phosphorylated eIF2α after 96 h (Figure 2A), whilst the B184 not expressing α-amylase showed eIF2α phosphorylation at the 48 h timepoint only (Figure 2A). AACK showed none or only minor phosphorylation of eIF2α at 48 h or 96 h, in agreement with its redox profile (Figure 2A, 1B). These results indicate that the increased phosphorylation of eIF2α in B184 is most likely linked to these strains increased capacity for α-amylase production (Figure 2A).

**Figure 2.**
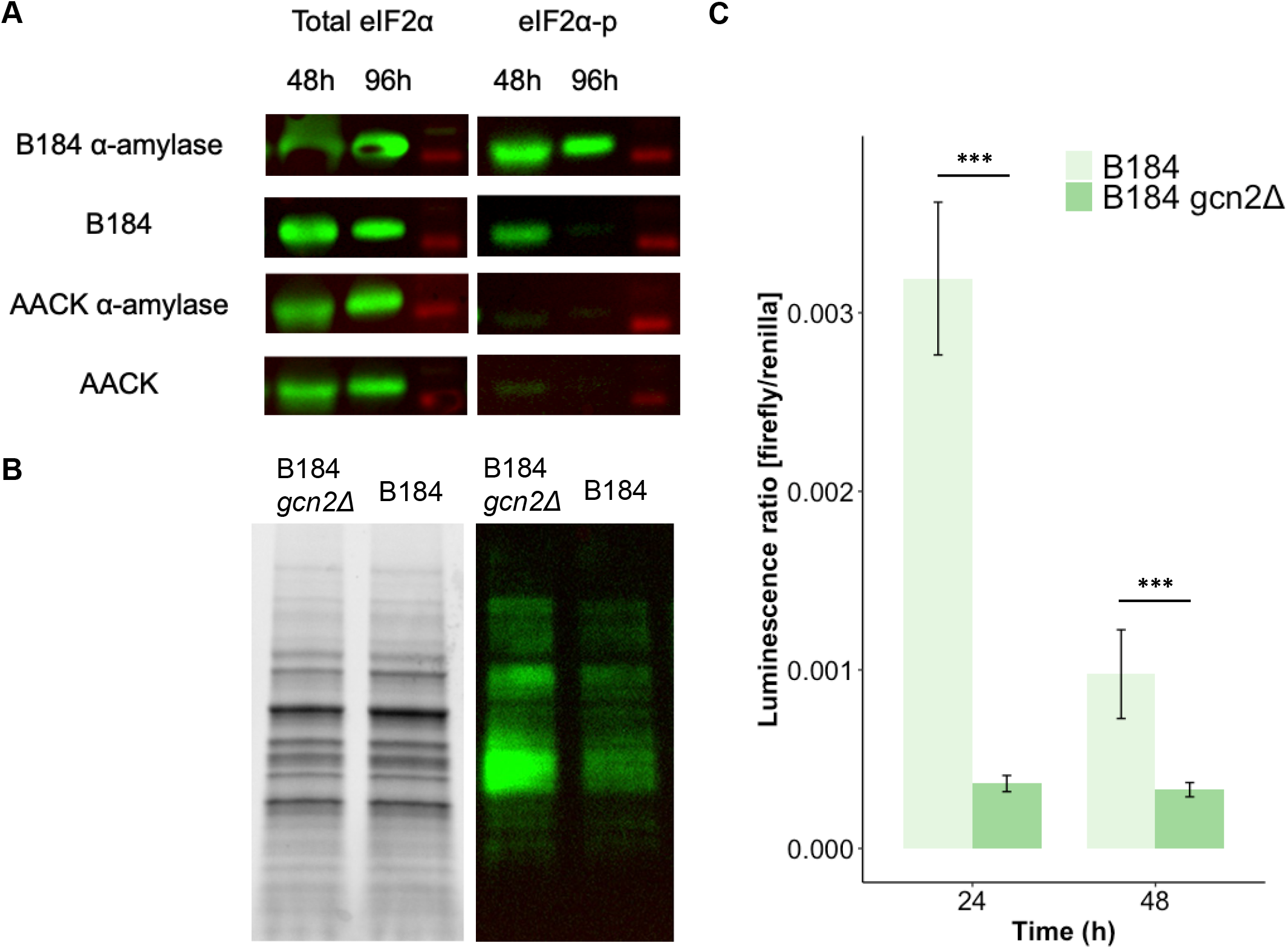
The protein kinase Gcn2 is active in the high-level production strain B184 under α-amylase expressing conditions. Upon removal of the *GCN2* kinase *GCN4* expression is reduced and overall translation is increased. **(A)** Western blot of total elF2α and eIF2α-phosphorylated. **(B)** Reducing SDS-page and Western blot of B184 *gcn2Δ* and B184 with primary antibody against puromycin during the exponential growth phase (OD=1). **(C)** *GCN4* expression assay based on a Firefly Renilla Luciferase Assay. The firefly luciferase gene is expressed under the control of the *GCN4* promoter and the renilla luciferase gene is under the control of the constitutive *PGK1* promoter. The luminescence ratio of firefly luciferase / renilla luciferase represents the normalized *GCN4* expression. *GCN4* expression levels in B184 (light green) and B184 *gcn2Δ* (green) after 24 and 48 h. Significance was determined using t-test with equal sample variance. Data are based on three biological replicates. * indicates P>0.05, ** indicates P>0.01 and *** indicates P>0.005 and the errors bars show the standard deviation.

### The removal of the GCN2 kinase leads to elevated rate of translation and decreased GCN4 expression

So far, our results indicate Gcn2 protein kinase activity in B184 producing recombinant proteins. To explore this further, we deleted *GCN2* in this strain and monitored how this would affect its best-known downstream targets, namely genes involved in general translation and the translation of the transcription factor *GCN4*. The rate of translation was measured using puromycin, a structural analog of aminoacyl-tRNAs that can be incorporated into the polypeptide chain but which prohibits further elongation (36). We included B184 and B184 *gcn2Δ* while producing α-amylase. Increased levels of puromycin-bound protein could be clearly seen in B184 *gcn2Δ* producing recombinant α-amylase compared to B184 when *GCN2* is expressed, suggesting that a higher rate of translation can be achieved when *GCN2* is absent (Figure 2B).

Next, we quantified the expression of *GCN4* which, alongside the general translation rate, is an indicator of Gcn2 activity. Several conditions activate Gcn2-mediated induction of *GCN4*, most of which are starvation related (18, 37, 38). Under non-starvation conditions, *GCN4* expression is inhibited through a post-transcriptional mechanism involving four uORFs that are preferentially translated over the *GCN4* ORF (38, 39). In contrast, during starvation and Gcn2 activation, the low levels of ternary complexes between eIF2-GTP and the initiator tRNA-Met, delay pairing with the AUG start codon sufficiently to bypass the uORFs and instead stimulate *GCN4* translation (38, 39). The expression of *GCN4* was determined using a luciferase assay with one construct expressing firefly luciferase under the control of the *GCN4* promoter and post-transcriptional regulatory regions, and a control renilla luciferase under the control of a constitutive promoter (40). We verified the functionality of the construct using chemically induced amino acid starvation (3-aminotriazole, Figure S5). The removal of the protein kinase Gcn2 in B184 producing recombinant α-amylase reduced *GCN4* expression significantly, in agreement with Gcn2 being the major activator of *GCN4* (Figure 2C) (41). In B184 cells, *GCN4* expression was visible at 24 h, however, its levels decreased at time-points during which Gcn2 activity increased (from 24 h to 48 h, Figure 2C). Taken together, these results show that in B184 producing recombinant α-amylase, the protein kinase Gcn2 is active in reducing both overall translation whereas the expression of Gcn4, in contrast, is reduced.

### The removal of the protein kinase Gcn2 leads to an improvement of recombinant α-amylase production in two engineered production strains

Having confirmed the activity of the protein kinase Gcn2 in B184 we wanted to quantify its impact on recombinant α-amylase production. We removed *GCN2* in two additional strains, AACK, as well as another strain, K17, which is optimized for α-amylase production and secretion by targeted engineering (5). K17 like B184 is engineered to improve protein production and reaches 5-fold α-amylase titers in bioreactors compared to the AACK strain (5, 35). Using these three strains both with and without α-amylase production, we quantified the amount of α-amylase produced, selecting time-points that reflected the different stages of growth. We cultivated the *gcn2Δ* and the control strains expressing recombinant α-amylase, for 96 h and sampled α-amylase after 24 h, 48 h, and 96 h. These results showed that final α-amylase titer in the media increased by approx. 2-fold in B184 upon *GCN2* removal (Figure 3A). Due to its previous engineering, B184 is already acknowledged as an efficient recombinant protein producer, particularly in combination with the CPOT expression plasmid [38], [39]. In comparison, for K17 *gcn2Δ*, the α-amylase titer increased 30%. The removal of the protein kinase Gcn2 also showed to have the highest impact on α-amylase production for all strains measured between 48 h and 96 h of cultivation (Figure 3A). Finally, for AACK, the removal of the protein kinase Gcn2 had no impact α-amylase titer at any timepoint during the 96 h of cultivation (Figure 3A). In addition to α-amylase productivity, we observed a significant increase in dry weight for B184 *gcn2Δ* while producing recombinant α-amylase, in comparison to B184 *GCN2*, (Figure 3B) which agrees with this strain having a relatively higher translation rate (Fig. 2B).

**Figure 3.**
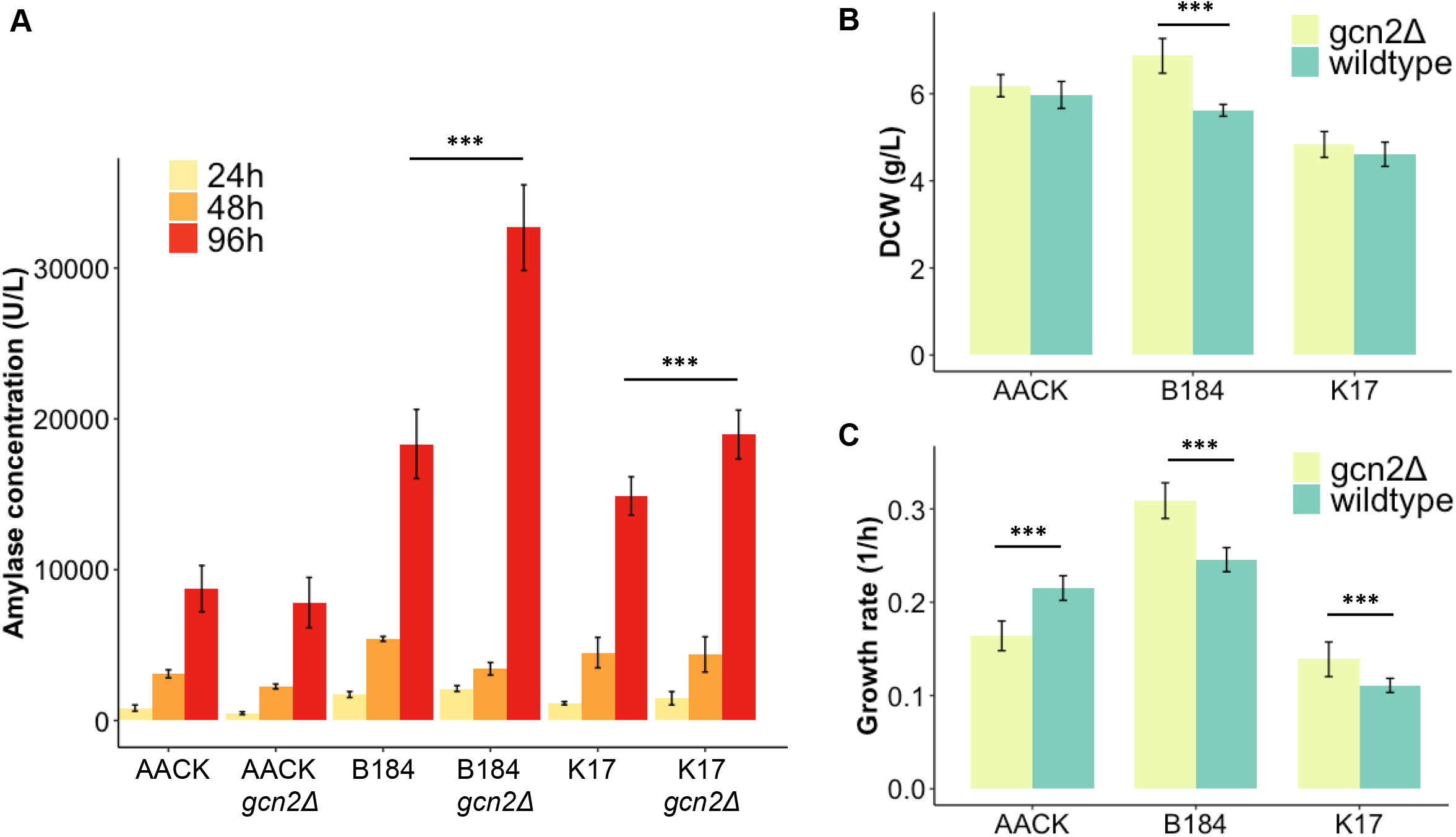
Removal of the protein kinase Gcn2 increases the α-amylase titer and improves growth parameters in two engineered high-level protein production strains. **(A)** α-Amylase concentration in the media after 24 (yellow), 48 (orange) and 96 (red) h of cultivations indicated with enzymatic assay, Data are the average of three biological replicates and two technical replicates. The significance is for the samples at 96 h. **(B)** Dry weight measurements after 96 h of cultivation in 24-well plates with the strains with intact *GCN2* (light green) and *GCN2* removed (green). Data is the average of three biological replicas and two technical replicas. **(C)** Exponential growth rates in 96-well plates with the strains with intact *GCN2* (light green) and *GCN2* removed (green). Data are the average of three biological replicates and three technical replicates. Significance was determined using t-test with equal sample variance. * indicates P>0.05, ** indicates P>0.01 and *** indicates P>0.005 and the errors bars show the standard deviation.

Lastly, we determined the exponential growth rates for all three strains with and without *gcn2Δ*. Here, growth rates significant increased for B184 *gcn2Δ* and K17 *gcn2Δ* whilst a decrease was observed for AACK *gcn2Δ* (Figure 3C). Therefore despite Gcn2 appearing to be beneficial for growth in AACK, for engineered strains wherein recombinant protein production is optimized for, this protein kinase has instead a detrimental impact. This supports our previous findings that *GCN2* is active for longer in engineered B184 strains, most likely due to its response to increased ROS levels during amylase production (Figure 2A & B, Figure 1A).

### The removal of the GCN2 kinase leads to decreased UPR activation whereas PDI1 expression is upregulated

To understand how *GCN2* may be linked to ROS production, we continued this study by examining the unfolded protein response (UPR) and the oxidative stress response, since these two mechanisms are intricately interconnected and have been previously linked to the control of translation (43). The UPR response in *S. cerevisiae* is activated by the Hac1 transcription factor, which itself is post-transcriptionally controlled by a splicing mechanism induced upon ER stress. Here, the spliced mRNA of *HAC1*, when translated into its active form leads to it inducing the transcription of the UPR response genes (13). We therefore measured the degree of *HAC1* mRNA splicing by qPCR to decipher if the UPR was being activated for our different strains. Interestingly, both B184 and B184 *gcn2Δ* strains, showed an increase in the *HAC1*^spliced^/*HAC1*^unspliced^ mRNA ratio from 24 h to 48 h suggesting that *HAC1* is more active in later stages of cell growth. When comparing B184 *gcn2Δ* to B184 however, the ratio of *HAC1*^spliced^/*HAC1*^unspliced^ mRNA decreased, both after 24 h and 48 h (Figure 4A), suggesting this strain experiences less ER stress during α-amylase production as a result of *GCN2* deletion.

We next selected several transcriptional Hac1 targets to check for their expression levels following *GCN2* deletion (Figure 4B). Here as anticipated we found that almost all genes had decreased expression, upon *GCN2* deletion suggesting the UPR was relatively inactive in these strains. The only exception, however, was *PDI1* which transcript increased 7-fold after 48 h in the B184 *gcn2Δ* strain. The expression of *PDI1’s* counterpart in disulphide formation, *ERO1*, was only modestly increased however (Figure 4B). The higher abundance of the *PDI1* transcript in B184 *gcn2Δ*, therefore seems independent of the UPR. The other known UPR target genes *KAR2, JEM1, EUG1, SCJ1*, and *LHS1* (Figure 4B) showed an expression similar to *ERO1*, in which their expression in B184 *gcn2Δ* was quite similar to in B184. The exceptions were *KAR2* and *JEM1* (Figure 4B). *KAR2* and *JEM1* showed a decrease in expression level which correlates with the lower level of *HAC1* splicing (Figure 4A).

**Figure 4.**
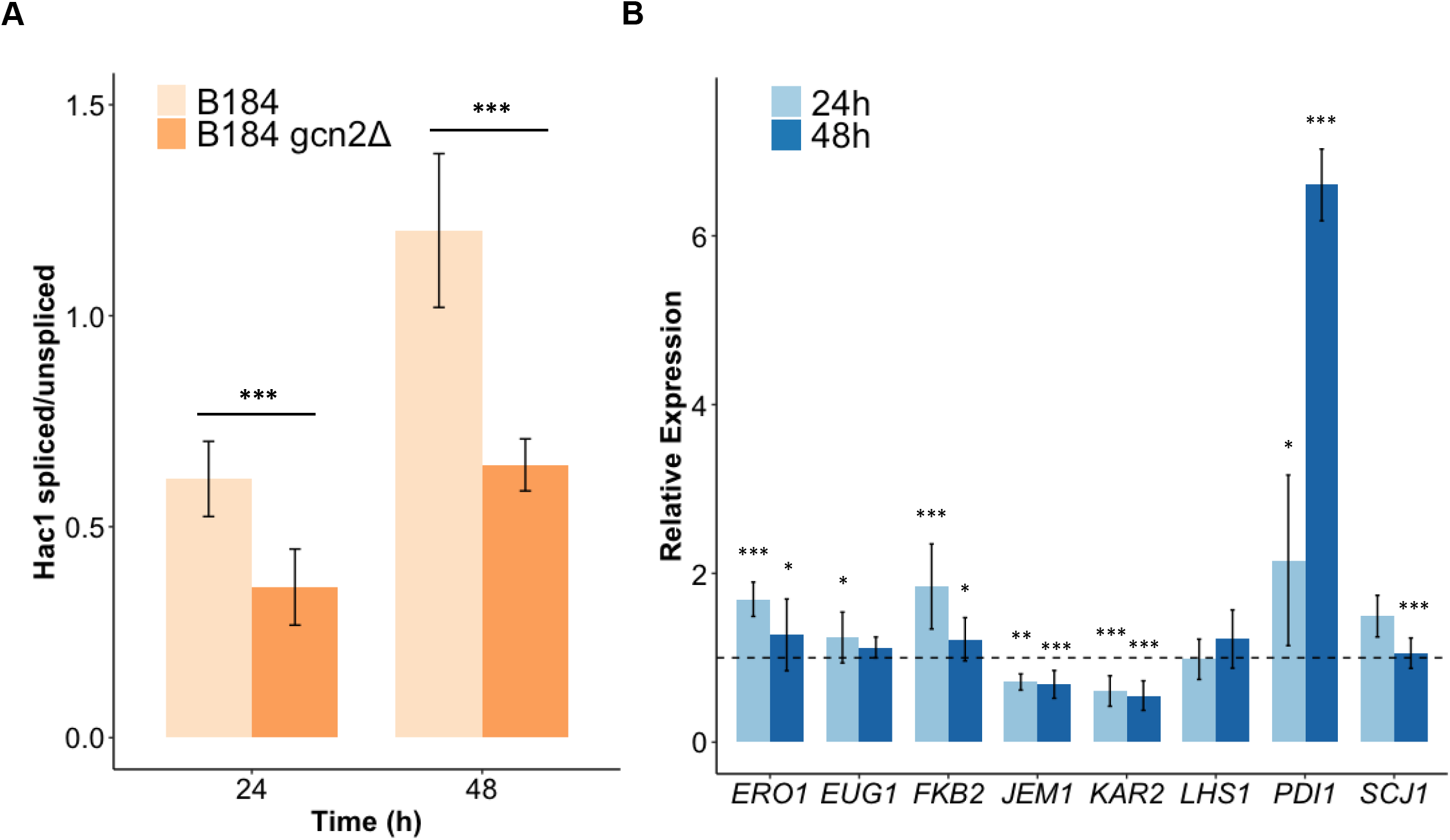
Removal of the eIF2 kinase Gcn2 reduces *HAC1* mRNA splicing while the expression of *PDI1* is strongly increased. For all the mRNA samples 3 biological and 3 technical replicates were included. **(A)** Ratio of spliced / unspliced HAC1 mRNA. The ratio is determined per sample based on the ΔCt of the spliced and the ΔCt of the unspliced HAC1 mRNA. It shows the Hac1 splicing of B184 (light orange) and B184 *gcn2Δ* (dark orange) after 24 h and 48 h. Significance is determined by difference of the ratios of the splicing between the two strains. Data are based on three biological with three technical replicates. Significance was determined using t-test with equal sample variance. **(B)** Expression levels of mRNA of UPR determined by qPCR. The data are analyzed using the ΔΔCt method and the data points indicate the relative expression of the genes encoding UPR-target proteins in B184 *gcn2Δ* compared to B184. The dashed line visualizes 1. For all the genes the mRNA was analyzed at 24 h (light blue) and at 48 h (dark blue). Significance is determined by the difference of the ΔCt per gene between the two strains. * indicates P>0.05, ** indicates P>0.01 and *** indicates P>0.005 and the errors bars show the standard deviation.

### Removal of the protein kinase Gcn2 leads to reduced H_2_O_2_ levels and an upregulation of antioxidant protein expression

So far our results suggest Gcn2 reduces ER stress during α-amylase production. As it is highly likely oxidative stress contributes to overall ER stress due to increased H_2_O_2_ levels during recombinant protein production, we next investigated the impact of Gcn2 on H_2_O_2_ production. We used the same setup as before with the Biolector and the roGFP2-Prx1 sensor to measure H_2_O_2_ levels in B184 *gcn2Δ* with and without recombinant α-amylase production. We added the data of the B184 *gcn2Δ* expressing α-amylase and the control to the plots shown in Figure 1A to provide the overview (Figure 5A). Across the duration of the entire cultivation the H_2_O_2_ levels were comparatively higher relative to B184 engineered for recombinant α-amylase production with *GCN2* intact (Figure 5A). In the control without recombinant α-amylase production the removal of *GCN2* does not impact the (Ox/Red) ratio during the cultivation. B184 *gcn2Δ* shows a (Ox/Red) ratio profile more similar to the controls. Considering that B184 *gcn2Δ* can achieve significantly higher amylase titers than when *GCN2* is expressed (Figure 3A), it is possible the concomitant lower H_2_O_2_ levels we observe is reflecting increased protein production in the ER, without the ER stress response being triggered by *GCN2*.

**Figure 5.**
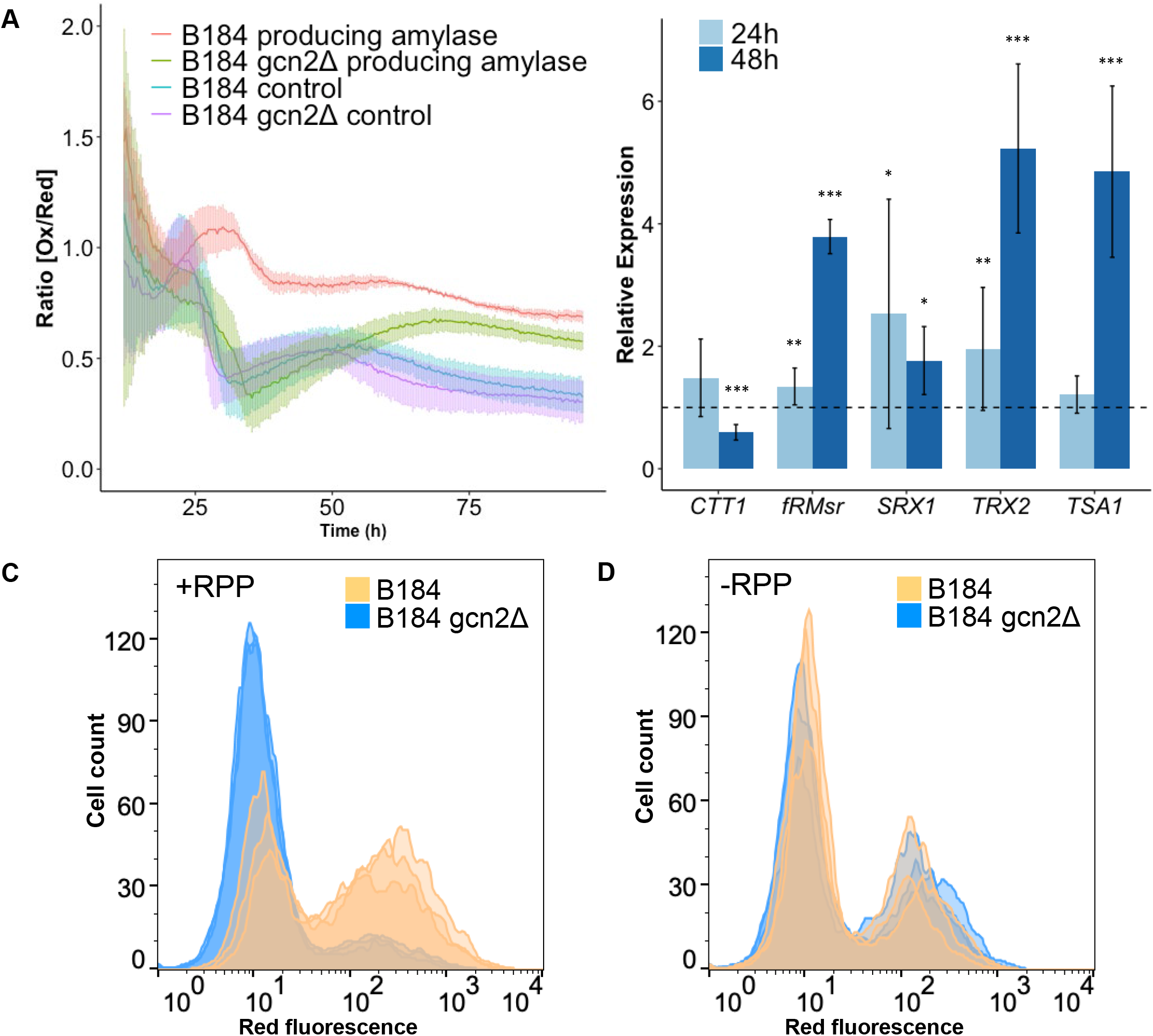
Removal of the protein kinase Gcn2 reduces H_2_O_2_ levels in B184 producing α-amylase, increases mRNA abundance of several antioxidant proteins and improves long time survival. **(A)** (Ox/Red) ratios over 96 h of cultivation for B184 measured with plasmid based *roGFP-PRX1*. B184 expressing α-amylase (red), B184 *gcn2Δ* expressing α-amylase (green), B184 without expressing α-amylase (blue) and B184 *gcn2Δ* without expressing α-amylase (purple). The light bars represent the standard deviations of three biological replicates and two technical replicates. The first 15 h were excluded due to too low signal. **(B)** Expression levels of mRNA of UPR determined by qPCR. The data were analyzed using the ΔΔCt method and the data points indicate the relative expression of the genes encoding anti-oxidant proteins in B184 *gcn2Δ* compared to B184. The dashed line visualizes 1. For all the genes the mRNA was analyzed at 24 h (light blue) and at 48 h (dark blue). Significance was determined by the difference of the ΔCt per gene between the two strains. Data are based on three biological with three technical replicates. Significance was determined using t-test with equal sample variance. * indicates P>0.05, ** indicates P>0.01 and *** indicates P>0.005. Survival measured with PI staining in combination with flow cytometry after 13 days of cultivation. Flow cytometry histograms with B184 (orange) and B184 *gcn2Δ* (blue) expressing recombinant α-amylase **(C)** and without recombinant protein production **(D)**, the figures contain three biological replicas per strain.

Among our B184 strains, growth profiles with the roGFP2-Prx1 sensor and the control plasmid without the sensor were comparable (Figure S6), highlighting that the inclusion of this sensor does not introduce any confounding effects in our analysis. In order to evaluate to what extent the decreased H_2_O_2_ levels observed in *gcn2Δ* cells reflected altered antioxidant levels, we next determined the expression of antioxidant proteins by qPCR. Except for *CTT1* a clear increase in relative expression levels could be seen for all anti-oxidant related genes tested, especially after 48 h when comparing B184 *gcn2Δ* to B184 (Figure 5B). *SRX1, fRMsr, TRX2*, and *TSA1* all showed elevated expression levels in B184 *gcn2Δ* compared to B184. The upregulation of most of the antioxidant genes we tested in B184 *gcn2Δ*, also correlates with this strain having lower overall levels of H_2_O_2_ (Figure S7). Taken together with results for B184 *GCN2* these results suggests that the presence of the protein kinase Gcn2 reduces the wildtype oxidative stress response.

### The removal of the Gcn2 kinase increases survival in recombinant a-amylase producing B184

ER stress has previously been suggested to increase the levels of mitochondrially-derived ROS, exerting a negative effect on cell survival (28). We therefore tested if the removal of *GCN2* with and without recombinant α-amylase production affected survival as a consequence of its impact on ER regulated UPR (Figure 4), H_2_O_2_ levels (Figure 5A), and on antioxidant gene expression (Figure 5B) in the cell. Using propidium iodide (PI) staining in combination with flow cytometry we could visualize and quantify the proportion of dead cells in our strain cell populations, whereby high fluorescence sub-populations represent dead cells and low fluorescence subpopulations indicate living cells. All strains showed 100% viability during the first 96 h of cultivation (Figure S7), after 13 days, however, the fraction of surviving cells increased in the B184 *gcn2Δ* cultures upon recombinant α-amylase production compared to B184 (Figure 5C) but not in the control without recombinant protein production (Figure 5D).

## Discussion

This work examined the roles of oxidants on recombinant protein production in yeast. We provide evidence for the accumulation of cytosolic H_2_O_2_ in cells engineered to produce high levels of α-amylase preferentially during the diauxic shift and post-diauxic shift growth phases. These are time-points during which amylase production peaks, suggesting that increased H_2_O_2_ is indeed a result of recombinant protein production. (6, 26)

Interestingly, a recent study found that increased endogenous H_2_O_2_ levels preferentially reacts with cysteines in proteins of the protein synthesis machinery, potentially explaining its inhibitory effect on protein production (44). Furthermore, H_2_O_2_ has been shown to repress protein synthesis in part through activating the eIF2α kinase Gcn2 (21). In agreement with these studies, we found that the protein kinase Gcn2 was activated in engineered *S. cerevisiae* strains producing recombinant α-amylase, downregulating translation and reducing α-amylase production (Figure 2A, 2B & 3A). These data are consistent with a model in which H_2_O_2_, accumulating as a result of recombinant protein production and secretion, represses cytosolic translation via the translation initiation factor (eIF2) kinase Gcn2 (Figure 6). In support for this model, the phosphorylation of eIF2 increases in a Gcn2-dependent manner upon α-amylase production (Figure 2A). Furthermore, cytosolic translation is maintained to a higher degree in Gcn2-deficient cells producing amylase (Figure 2B). Unexpectedly, however, we found that both the ER specific UPR and oxidative stress responses were affected by the removal of *GCN2*. Whereas the UPR decreased in cells lacking Gcn2 (Figure 4B), the antioxidant response was increased (Figure 5B) correlating with decreased cytosolic H_2_O_2_ (Figure S7).

**Figure 6.**
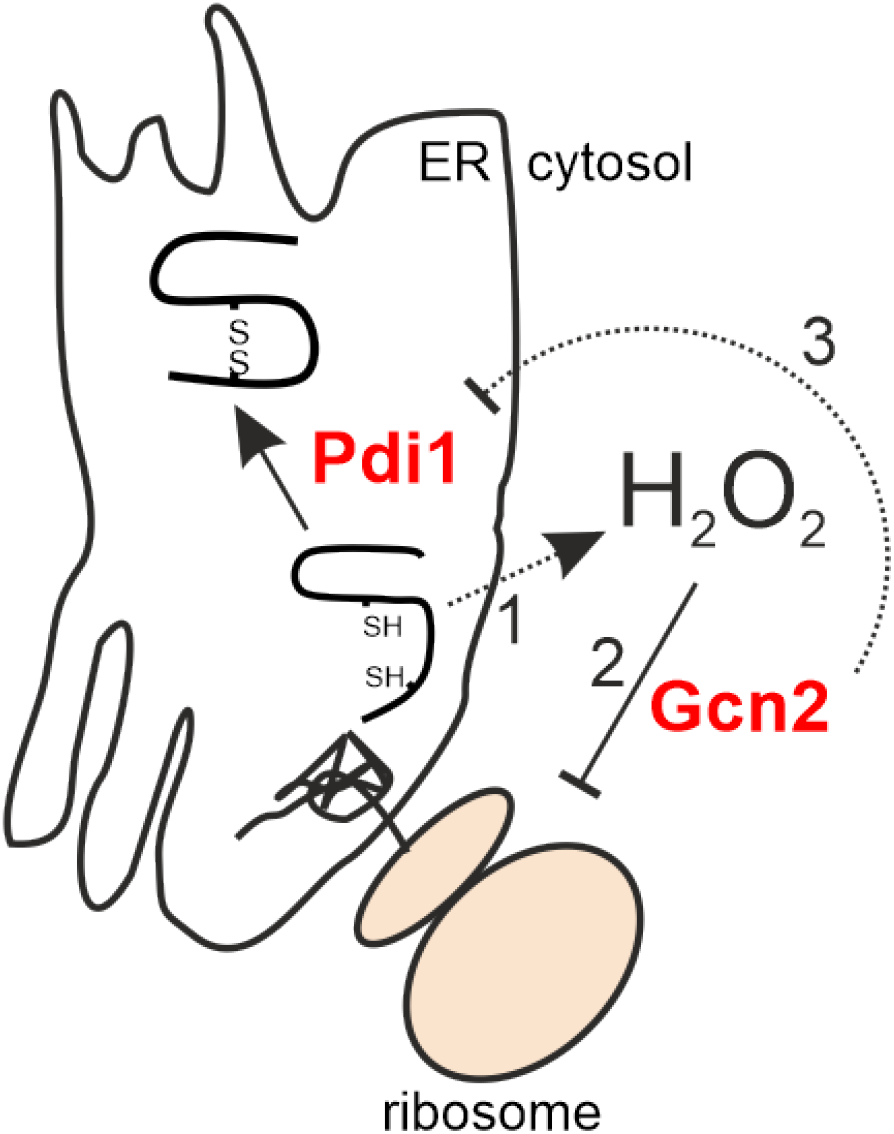
Model of mechanisms by which Gcn2 affects protein synthesis and ER oxidative folding. Recombinant protein production leads to the accumulation of H_2_O_2_ in the cytosol via an unknown mechanism (1). H_2_O_2_ activates the translation initiation factor (eIF2) kinase Gcn2 (2) causing the repression of protein synthesis. Through an unclear mechanism Gcn2 also appears to exert an inhibitory effect on both anti-oxidant expression and *PDI1* transcription (3), suggesting that cytosolic protein synthesis is coordinated with ER oxidative folding.

### Reduction of the UPR in B184 gcn2Δ

The UPR has previously been coupled to elevated H_2_O_2_ levels and oxidative stress. In this work, Haynes et al. observed that in ERAD-deficient cells challenged with increased levels of misfolded proteins, the removal of the UPR reduced oxidative stress and improved fitness (28). We observed a decrease in the UPR and reduced H_2_O_2_ levels upon loss of Gcn2. The level of oxidative stress has previously been thought to be result of folding in the ER (11, 12). This is not coherent with our data, however, since we also observe an increased α-amylase production upon Gcn2 removal (Figure 3A). Besides, the UPR target genes show a variable expression pattern.

A somewhat surprising finding in this study was the rather strong induction of *PDI1* (Figure 4B) that appears to be unrelated to the UPR. In particular, we observe an almost 7-fold induction of the *PDI1* transcript in B184 cells lacking Gcn2 (Figure 4B). Previous studies have shown overexpression of *PDI1* to have a positive influence on protein production, e.g. of α-amylase (5, 34). This indirectly induced overexpression of *PDI1*, caused by the absence of protein kinase Gcn2, could thus be an additional explanation for the increase in α-amylase production in this strain. The strain B184 indeed carries a chromosomal duplication leading to two copies of the *PDI1* gene in the genome (34). The mechanism that results in this strong induction of *PDI1* in the absence of Gcn2 is, presently, unknown. Two independent large-scale transcriptomic studies, however, point out the transcriptional activator of ribosomal genes, Sfp1, as a regulator of *PDI1* (45, 46), suggesting coordination between the cytosolic protein synthesis machinery and ER localized oxidative folding (Figure 6).

The Hac1-mediated induction of the UPR occurs via binding to UPR response elements, UPREs. Previous research has shown that there are at least three different UPREs, with the expression of associated target genes being dependent not only on Hac1 activity, but also by Gcn4 expression, the downstream target of Gcn2 (47). It has also been shown that the removal of protein kinase Gcn2, blocks the expression of UPR genes independently of *HAC1* splicing upon oxidative folding stress (47). Other studies indicate however that Hac1 binds independently of other factors to at least two of the UPREs (48). Specifically, *KAR2* contains the UPRE referred to as UPRE-1 in its promotor (47) and so does the promotor of *JEM1* in the strain we used. Their downregulation is thus coherent with the reduced *HAC1* mRNA splicing observed in B184 *gcn2Δ* (Figure 4A). Based on our results, the expression of *KAR2* and *JEM1* correlate with the *HAC1* mRNA splicing ratio indicating that the UPRE-1 mediated expression of those genes is neither influenced by Gcn2 nor Gcn4 activity.

### Removal of the Gcn2 kinase and its impact on H_2_O_2_ levels

Interestingly, we could demonstrate that the removal of the protein kinase Gcn2 in B184 leads to a decrease in intracellular H_2_O_2_ levels, even though α-amylase production is higher (Figure 5A, 1A, 3A). H_2_O_2_ is a byproduct of the iterative process of forming correct disulfide bridges in proteins (28, 49). One could therefore assume that more protein produced would lead to a higher level of H_2_O_2_. However, H_2_O_2_ levels in the ER may be maintained mostly independently of cytosolic H_2_O_2_ levels (50). In agreement with our data suggesting that H_2_O_2_ is potentially originating outside of the ER, while still interfering with ER oxidative homeostasis, mitochondrially-derived H_2_O_2_ has been shown to increase cytosolic H_2_O_2_ levels in ERAD deficient cells (28).

We find also that reduced levels of cytosolic H_2_O_2_ in cells lacking Gcn2 correlate with the upregulation of several anti-oxidant genes such as *TSA1, TRX2, SRX1* and *fRMsr* (Figure 5A & 5B). Trx2 is a thioredoxin and is known to reduce cytosolic 2-Cys peroxiredoxins like Tsa1, whilst Srx1 is a sulfiredoxin that reactivates hyperoxidized Tsa1 (51). Interestingly, in support for an importance of Gcn2 in the anti-oxidant response, this protein has previously been shown to be required for high-level translation of the *SRX1* mRNA (51). Furthermore, *TSA1, TRX2*, and *SRX1* are all known targets of Yap1, a transcription factor that responds to elevated H_2_O_2_ levels (52–54). These genes’ increased expression therefore suggest that Yap1 may be activated in B184 *gcn2Δ* while producing recombinant α-amylase. Previous work by Delic et al, showed that by overexpressing *YAP1* the redox balance of the cytosol in a recombinant protein producing *P. pastoris* strain was restored (55).

With the findings in this study, we conclude that in two strains engineered for optimized protein production, the protein kinase Gcn2 is responsible for mediating a negative feedback loop affecting both cytosolic translation and the secretory pathway. By removing this H_2_O_2_-mediated feedback loop recombinant protein production is improved, indicating that the reduction of translation via endogenous oxidants can limit the productivity of yeast cells. The active protein kinase Gcn2 negatively affects several processes in the cell including ER stress and H_2_O_2_ levels. Such findings are relevant for the engineering of production hosts for biotechnological production processes but also in basic research through the understanding of feedback loops present in multiple biological systems.

## Materials and Methods

### Strains and plasmids

Three previously constructed *S. cerevisiae* strains were used in this study. CEN.PK 530.1CK *[MATα URA3 HIS3 LAU2 TRP1 SUC2 MAL2-8^c^ tpi1*(41-707)] further referred to as AACK. Previous studies have engineered AACK to improve protein production leading to two strains, B184 and K17 (5, 34). B184 is generated by UV mutagenesis and K17 has the following genotype AACK *[Δhda2 Δvps5 Δtda3 PGK1p-COG5 Δgos1:: amdSYM-TEF1p-PDl1]*, AACK, B184, and K17 additionally have a disrupted *TPI1* gene. To complement this deficiency, we use the pAlphaAmyCPOT plasmid with an expression cassette for α-amylase. This cassette has a α-leader sequence and an α-amylase gene from *Aspergillus oryzae* (42). As a control, an empty CPOT plasmid was used. The *GCN2* gene was disrupted with help of plasmid pECAS9-gRNA-kanMX which contains both a cas9 gene and a gRNA expression cassette (56). The plasmids pECAS9-gRNA-kanMX-GCN2 and pECAS9-gRNA-kanMX-URA3 were made using the pECAS9-gRNA-kanMX-tHFD1 as the template (56). First, the backbone was obtained by linearizing pECAS9-gRNA-kanMX-tHFD1 by digestion with MunI and EcoRI. The ‘left’ fragment was constructed with primer #54 in combination with either #53 *(GCN2)* or #61 *(URA3)* and the ‘right’ fragment was constructed with primer #55 in combination with either #52 *(GCN2)* or #60 (*URA3*). The correct assembly of the plasmids was confirmed by sequencing using primer #42. The genomic deletion was verified using primers pairs #38 and #39 for *GCN2* and #40 and #41 for *URA3*. The gRNA, repair fragments, and verification primers can be found in the Supplementary data for *GCN2* and *URA3* genes (Table S1). The plasmids used in this study can be found in Table S3. *E. coli* DH5α was used for plasmid amplification.

### Media and culture conditions

Media used for *S. cerevisiae* strain construction were YPD, YPE, YPEG, SD-URA. The experiments were always performed at 30 °C and 220 rpm. YPD medium contained 10 g/L yeast extract, 20 g/L peptone, and 20 g/L glucose and was used for all cultures unless otherwise mentioned. For the selection of the kanMX marker on the CRISPR plasmid, 200 mg/L G418 (Formedium, Hunstanton, UK) was added to the YPD medium. The YPE medium contained 10 g/L yeast extract, 20 g/L peptone, 20 g/L absolute ethanol and was solely used as a solid medium. For liquid cultivations 30 g/L glycerol was added to YPE and the medium was referred to as YPEG. Both YPE and YPEG were only used for *S. cerevisiae* strains without CPOT plasmids since those are unable to ferment glucose as the sole carbon source (57). SD-URA contained 20 g/L glucose, 6.7 g/L yeast nitrogen base without amino acids, and 0.77 g/L complete supplement mixture without uracil (CSM-URA, Formedium) This medium was only used to verify the deletion of the *URA3* gene. To solidify media 20 g/L agar (Merck Millipore) was added. The protein expression and physiological experiments were performed in SD2XSCAA media with glutamine instead of glutamate. SD-2XSCAA medium contained 20 g/L glucose, 6.9 g/L yeast nitrogen base without amino acids, 190 mg/L Arg, 400 mg/L Asp, 1260 mg/L Gln, 130 mg/L Gly, 140 mg/L His, 290 mg/L Ile, 400 mg/L Leu, 440 mg/L Lys, 108 mg/L Met, 200 mg/L Phe, 220 mg/L Thr, 40 mg/L Trp, 52 mg/L Tyr, 380 mg/L Val, 1 g/L BSA, 5.4 g/L Na_2_HPO_4_, and 8.56 g/L NaH_2_PO_4_·H_2_O and had a pH of 6.4.

Characterization of the roGFP2 sensors was performed in Delft synthetic medium (58) and the verification of the luciferase expression in defined synthetic medium lacking uracil, using 14 mL cultivation tubes (59). Protein production experiments and *GCN4* expression experiments were performed at 30°C at 220 rpm in aerated 24-wells plates CR1224 (Bioscreen) with a volume of 2.5 mL and a start OD_600_ of 0.01. All other experiments were grown in 100 mL shake flasks with 10 mL SD2xSCAA medium and a starting OD_600_ of 0.01. The cultures for qPCR analysis were grown in a volume of 20 mL with a starting OD_600_ of 0.01. *E. coli* cells were grown in Luria-Bertani (LB) media at 37°C and 200 rpm. Selection medium contained 80 mg/L Ampicillin. The transformation procedure used for *E.coli* was according to a known protocol (60).

### Molecular biology techniques

*S. cerevisiae* strains were transformed according to the protocol using the Li/Ac SS carrier method (61). 500 ng of DNA was used for the transformation of plasmids and an additional 1 μg repair fragment when required. To verify deletions or test for the presence of the CPOT plasmids colony PCR was performed using SapphireAmp fast PCR mix (TaKaRa Bio). For DNA construction, Phusion High Fidelity DNA polymerase (Thermo Scientific) was used. Restriction digestion was performed using FastDigest (Thermo Scientific) products. All techniques were used according to the manufactures protocols unless otherwise stated.

### α-Amylase assay

Cells were harvested after 24 h, 48 h, and 96 h respectively. Cells were pelleted by centrifugation at 4°C, 8000 rpm for 5 min, then the supernatant was used for the α-amylase quantification assay. The Ceralpha kit (Megazyme) was used with α-amylase from *Aspergillus oryzae* as the standard. The assay was performed according to the manufacture’s protocol with an exception on the preparation of buffer A. Since the protein was dissolved in the media, instead of preparing buffer A and dissolving solidified protein, we used a mixture of media and Milli Q water, depending on the concentration of α-amylase, to make buffer A with the correct concentration and protein. We used either a dilution of 200X or 400X depending on the concentration of α-amylase in the media.

### Growth profiler

The *S. cerevisiae* strains were cultivated for 48 h in 250 μL SD2xSCAA medium at 30°C and 1200 rpm in 96-well plates (Enzyscreen CR1496d). Growth curves were measured using a Growth Profiler 960 (Enzyscreen). Three independent colonies per strain were grown in 1 mL SD2XSCAA media in 7 mL cultivation tubes after an overnight culture. The cells were then inoculated in technical triplicates with a starting OD_600_ of 0.005.

### Microbioreactor cultures

*S. cerevisiae* strains were cultivated for 96 h in 1 mL SD2xSCAA media at 30 °C and 1200 rpm in ‘Flowerplates’ The characterization of the sensors was performed in Delft minimal media and the experiments in SD2xSCAA media. Three independent colonies per strain were grown in 1 mL SD2XSCAA media in 7 mL cultivation tubes after an overnight culture. Cells were then inoculated in technical duplicates with a starting OD_600_ of 0.005. For measuring the biomass, excitation and emission at 600 nm was used with gain 20, for the oxidation of cysteine, an excitation at 405 nm and emission at 520 nm with gain 100 was used and for the reduction of cysteine, an excitation at 488 nm and emission at 520 nm with gain 100. All wells were measured every 20 min by a Biolector microbioreactor system (M2p-Labs).

### Ox/Red ratio determination

Background fluorescence was determined using strains carrying an empty p416 vector. We used biological duplicates of these controls with technical duplicates. The natural fluorescence per strain was determined at both 405 nm (Ox) and 488 nm (Red). For both wavelengths, the average of the natural fluorescence was determined. These average values were subtracted from the Ox and Red measurements of all the separate replicates with the roGFP2 sensors. The GFP signals with the natural fluorescence subtracted were used to determine the Ox/Red ratio per replicate per strain. The final Ox/Red ratio was determined by taking the average of the ratios per strain. R Studio software was used for all data analysis (62).

### qPCR

Cells were harvested after 24 h and 48 h respectively, cells were then instantly cooled on ice and centrifuged at 4°C, 6000 rpm for 3 min. The supernatant was discarded, and the pellet was snap-frozen using liquid nitrogen. For RNA extraction, the RNeasy Kit (Qiagen) was used according to the manufacturers protocol. For cDNA synthesis, the Quantitect Reverse Transcriptase Kit (Qiagen) was used. For the qPCR, the DyNaMo ColorFlash SYBR Green qPCR Kit was used. All primers listed (Table S2) were verified using the MIQE guidelines with *ACT1* used as the reference gene.

### Puromycin treatment

Yeast cells were grown in SD2xSCAA and grown until the mid-exponential phase (OD_600_ ~1). Cells were then normalized to OD_600_ 1, then harvested and collected by centrifugation before being incubated in 100 mL PBS with 1 mM puromycin for 10 min at 30°C, 220 rpm. Cells were then collected by centrifugation, and intracellular proteins were extracted as described previously (63). 10 μL of the cell extracts were then used for SDS-page and Western Blot analysis.

### eIF2α protein extraction

For the intracellular protein extraction of the elongation factor eIF2α, protein extraction with LiAc/NaOH was performed as in (64). 5 OD_600_ of yeast cells were harvested after 24 h and 48 h or and 10 OD_600_ at 72 h and 96 h. 10 μL of the cell extracts was then used for SDS-page and Western Blot analysis.

### Western blotting

Samples and controls were loaded on and separated with Stain free 4-20% gels (Biorad). Proteins were transferred onto 0.45 micron PVDF membranes (Bio-rad) using the Trans-Blot Turbo transfer system (Bio-rad). The blot was blocked using Western blocker solution (Sigma Aldrich) and incubated in either anti-total eIF2α (1:1000) or anti-puromycin (1:1000), or eIF2α-phosphorylated (Ser-51; Invitrogen; 1:1000) followed by incubation with either anti-mouse (1:5000) or anti-rabbit (1:5000). Both secondary antibodies are HRP-conjugated and were visualized using West Pico Plus HRP substrate (Thermo Fischer) and measured with a ChemidoC XRS image analyzer (Bio-Rad).

### Viability measurements

Cell viability was measured using propidium iodide (Invitrogen) staining as described previously (63). Samples were taken after 1, 2, 3, 4, and 13 days of cultivation in 10 mL of SD2xSCAA media. Fluorescence was measured with a Guava easyCyte 8HT system (Merck Millipore). For each sample, 5000 cells were counted. The cultivations were performed in biological triplicate and unstained cells were used as a negative control for the fluorescence measurements.

### *GCN4* activity assay

The luciferase construct was tested in a *S. cerevisiae* BY4742 strain in which the pVW31 plasmid was transformed. Three biological replicas were cultivated in 7 mL cultivation tubes to which 10 mM (final concentration) 3-AT was added and incubated for 30 min. The luminescence was checked before and after the addition of 3-AT. For the *GCN4* expression experiment, cells were harvested after 24 h and 48 h. 2 mL of culture was centrifugated for 5 min at 35000 rpm at 4°C. The supernatant was then discarded, and cells washed in 1 mL cold water. Cells were resuspended in 300 μL PBS buffer with protease inhibitors and added to lysin matrix tubes (MP Bio). The mixture was Fast prepped at 5000 rpm for 20 sec 3 times with incubation of the samples on ice between runs. The mixture was then centrifuged for 10 min at max speed at 4°C and 100 μL of clear supernatant was harvested and stored at −20°C. Luminescence was measured with a FluoStar Omega plate reader (BMG Labtechnologies) and treated with the protocol and reagents of the Dual-Luciferase Reporter Assay System (Promega). All reagents were used accordingly to the manufactures protocol.

## Acknowledgments

The authors thank Dr. Xin Chen for the help with the flow cytometer and qPCR setup and operations.

